# Measuring the Importance of Vertices in the Weighted Human Disease Network

**DOI:** 10.1101/436402

**Authors:** Seyed Mehrzad Almasi, Ting Hu

## Abstract

Many human genetic disorders and diseases are known to be related to each other through frequently observed co-occurrences. Studying the correlations among multiple diseases provides an important avenue to better understand the common genetic background of diseases and to help develop new drugs that can treat multiple diseases. Meanwhile, network science has seen increasing applications on modeling complex biological systems, and can be a powerful tool to elucidate the correlations of multiple human diseases. In this article, known disease-gene associations were represented using a weighted bipartite network. We extracted a weighted human diseases network from such a bipartite network to show the correlations of diseases. Subsequently, we proposed a new centrality measurement for the weighted human disease network in order to quantify the importance of diseases. Using our centrality measurement to quantify the importance of vertices in the weighted human disease network, we were able to find a set of most central diseases. By investigating the 30 top diseases and their most correlated neighbors in the network, we identified disease linkages including known disease pairs and novel findings. Our research helps better understand the common genetic origin of human diseases and suggests top diseases that likely induce other related diseases.

## Author summary

## Introduction

During the past decades, significant progress has been made in our understanding of human diseases [1]. However, the genetic architectures of complex diseases are still largely unclear. Many common diseases tend to be related to each other, and it is suspected that they may share common genetic origin. Thus, studying the correlations of human diseases has the potentials of better understanding the genotype to phenotype mapping [2, 3] and better predicting disease association genes [4, 5, 6, 7, 8]. Furthermore, learning which diseases are correlated can help use existing drugs to treat multiple similar diseases [9, 10, 11, 12, 13].

Meanwhile, network science is a rising field where entities and their complex relationships are studied on a global scale [14, 15, 16], and has seen increasing applications to perform advanced analysis on biomedical data [17, 18, 19, 20, 21, 22]. There are various cellular components in the human body that interact with each other within the same cell or across different cells [15]. A network called the *human interactome* can be constructed according to the interactions of those different cellular components. Each component can be represented as a vertex in the network and interactions among them can be captured as links (or edges) connecting pairs of the cellular components. Those cellular components can be proteins or metabolites, and the network refers to protein-protein interaction (PPI) network [23, 24, 25] or metabolic network [26, 27, 28].

Some studies aimed at identifying the correlations among diseases through network analysis [15, 29, 30]. Goh *et al.* [31] constructed a human disease network (HDN) by connecting pairs of diseases when they share common association genes. Of 1,284 diseases in the HDN, 867 have at least one link to other diseases, and 516 form a giant component, suggesting that the genetic origins of most diseases, to some extent, are shared with other diseases. Moreover, the HDN naturally and visibly clustered according to major disease classes such as cancer cluster and neurological disease cluster. Zhou *et al.* [32] extracted over twenty million bibliographic records from PubMed [33] in order to obtain 147,978 connections between 322 symptoms and 4,219 diseases. A human symptoms-disease network (HSDN) was then constructed and was able to show the symptom similarity between all pairs of diseases (7,488,851 links) in the network. The weight of links represented the similarity of symptoms between two diseases. They showed that the correlations among diseases were significantly related to the genetic associations that each pair of diseases had in common as well as the interactions between their related proteins. Lee *et al.* [34] built a disease metabolism network in order to study disease comorbidity for better disease prediction and prevention. Two diseases are connected with each other if a mutated enzyme catalyzes metabolic reaction between them. Their results show that diseases with higher degrees, i.e., connecting with many other diseases, have a higher rate of prevalence and mortality.

Measuring the centrality of vertices helps identify important vertices in the network in terms of connecting to all other vertices. Centrality measures have been used frequently to analyze biological networks over the past decades [35, 36, 37]. The most common centrality measures include degree (the total number of neighbors), closeness (the total distance to all other vertices), and betweenness (the fraction of locating on the shortest paths of all pairs of vertices) [38]. Despite wide applications in biological networks, these centrality measures are rather general and may not be able to capture all the properties of vertices in the context of biological networks. Therefore, carefully tailored centrality measures are needed for specific network of interest, in this study, the human disease network.

Köhler *et al.* [39] proposed a vertex importance measure for disease genes in the context of PPI networks. They used a random walk strategy to assess the distance between vertices in the network, and reported improved performance comparing with conventional distance-based centrality measures. Wu *et al.* [40] integrated PPI networks with gene expression data in order to rank disease genes associated with various cancers. They showed that their method was able to find replicable high-rank genes using different datasets. Martinez *et al.* [41] proposed a generic vertex prioritization method using the idea of propagating information across data networks and measuring the correlation between the propagated values for a query and a target set of entities. The authors tested their method by ranking disease genes associated with Alzheimer’s disease, diabetes mellitus type 2 and breast cancer. They reported some new high-rank association genes that could bring new insights into the diseases.

In the article, we propose a new method for the construction of a weighted human disease network (WHDN) and a new centrality measure to identify the most important diseases. First we use a large database of disease-gene associations to build a weighted bipartite disease-gene network, and then construct a weighted disease network where link weights capture the strengths of the pairwise disease correlations. After the backbone extraction of the WHDN, we design a centrality measure for the context of the WHDN that considers not only the degree of a vertex but also the importance of its incident edges. Finally, we compare our new centrality measure with degree, closeness and betweenness by evaluating the network efficiency decline rate with the removal of top-ranked vertices by each centrality measurement.

## Methods and Results

Given the multiple-step pipeline structure of this study, we show the result of each step after the description of the corresponding method.

### Disease-Gene Associations (DGAs)

The data used in this project contains disease-gene associations (DGAs) from multiple curated databases including UNIPROT, CTD (human subset), PsyGeNET, Orphanet, and HPO. The disease-gene association data are conducted by DisGeNet group, available on DisGeNET v4.0 [42]. The current version of the data set contains 130,821 DGAs, between 13,075 diseases and 8,949 genes. Each DGA is assigned with a score 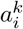, for disease *i* and gene *k*, within the range of [0,1] based on its level of evidence, the number and the type of database sources supporting the DGA, and the number of publications verifying the association between the gene and the disease [42]. We first clean up the data in order to ensure that all diseases and genes in the dataset are unique and that there is no replication of disease-gene associations. Next, since we would like to consider the correlation among all diseases, we keep diseases and syndromes in the dataset for our analysis and remove injuries or poisonings, anatomical abnormalities, acquired abnormalities, mental or behavioral dysfunctions, signs or symptoms, findings, congenital abnormalities, neoplastic processes, and pathologic functions. We use DisGeNet web-based application [42] for this filtering.

### Network Construction

#### Bipartite Disease-Gene Association Network

The best representation for depicting the associations among genes and diseases is a bipartite graph, which is called the disease-gene association network in this research. The bipartite graph contains two different sets of vertices. One set includes diseases and another one contains genes. By definition, no edge is allowed to connect a pair of vertices in the same set of vertices in a bipartite graph. That is, there can be no link either between a pair of diseases or a pair of genes. There is an edge between a gene and a disease if there is an association between them. Their link weight is assigned as the score 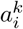, for disease *i* and gene *k*, computed in the DGA database described in the previous section. A sample subgraph of the bipartite network is shown in Figure 1.

Figure 2 depicts the degree distributions of diseases and genes in the bipartite disease-gene association network. For the set of diseases, the maximum degree is 564, of the disease *epilepsy,* and the average degree is 5.43. In Figure 2 a), the degree distribution of the diseases is right-skewed and approximately follows a heavy-tailed distribution, indicated by the straight linear fit on a log-log scale. For the set of genes, the maximum degree is 111, of the gene LMNA, and the average degree is 5.81.

The bipartite network is comprised of multiple connected components with a single giant component. Figure 3 shows its distribution of the size of connected components. The giant component has 10,212 vertices consisting of 5,278 diseases and 4,934 genes. Apart from the giant component, all other connected components are small with a size varying from two to nine, and most of them are only single pairs of one disease and one gene. Figure 3 shows that there is a considerable number of components with two vertices, i.e., 844 isolated disease-gene pairs. Since we are interested in investigating the large-scale genetic correlations of human diseases, we focus the giant component of the disease-gene bipartite network in the downstream analyses.

**Fig 1.**
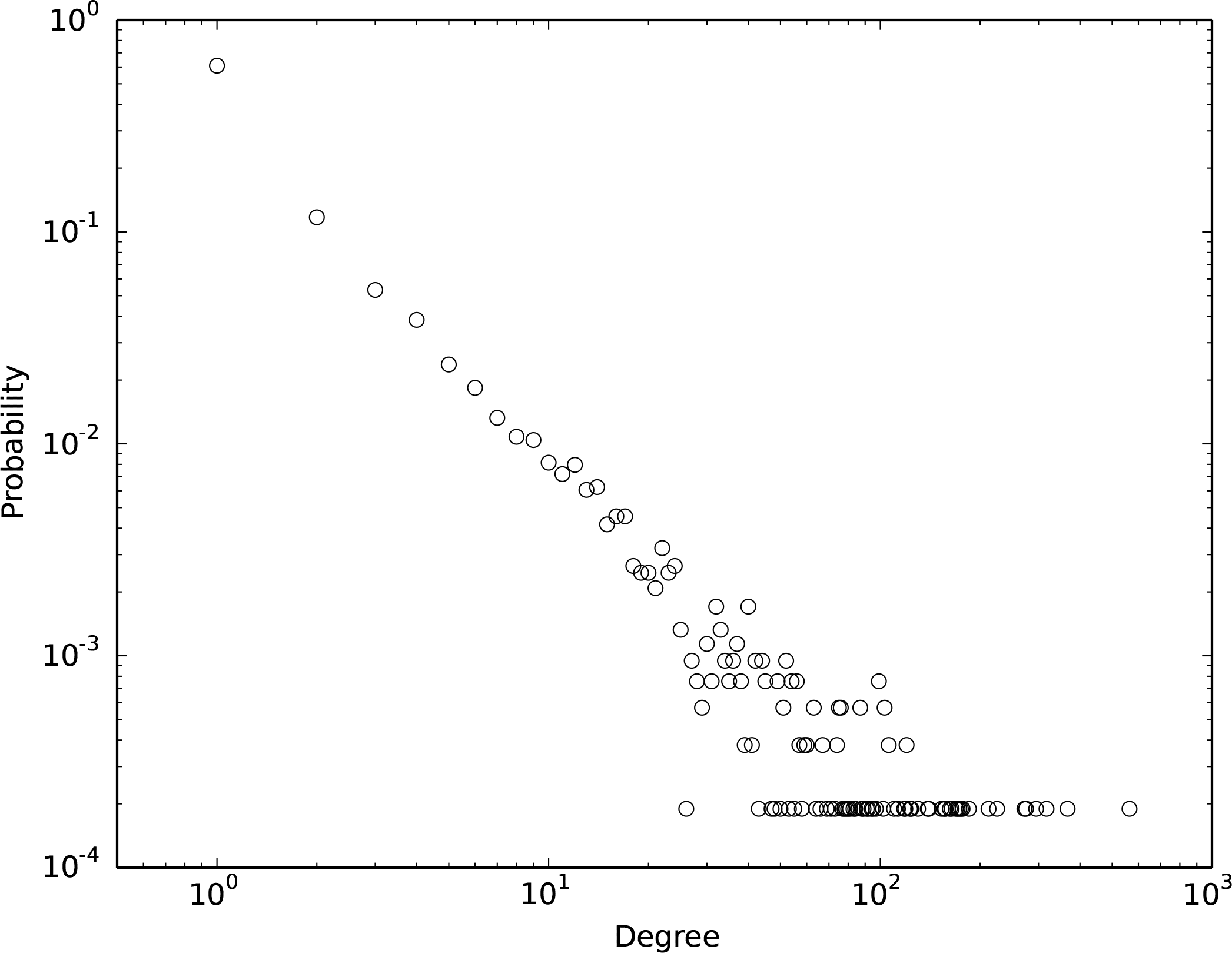
An example subgraph of the human disease-gene association network. The bipartite network has two sets of vertices, i.e., genes and diseases, represented by rectangle and gray ellipses respectively. An edge connects a disease and a gene if there is a known association between them. The weight of an edge indicates the strength of the DGA 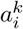 between disease *i* and gene *k*.

**Fig 2.**
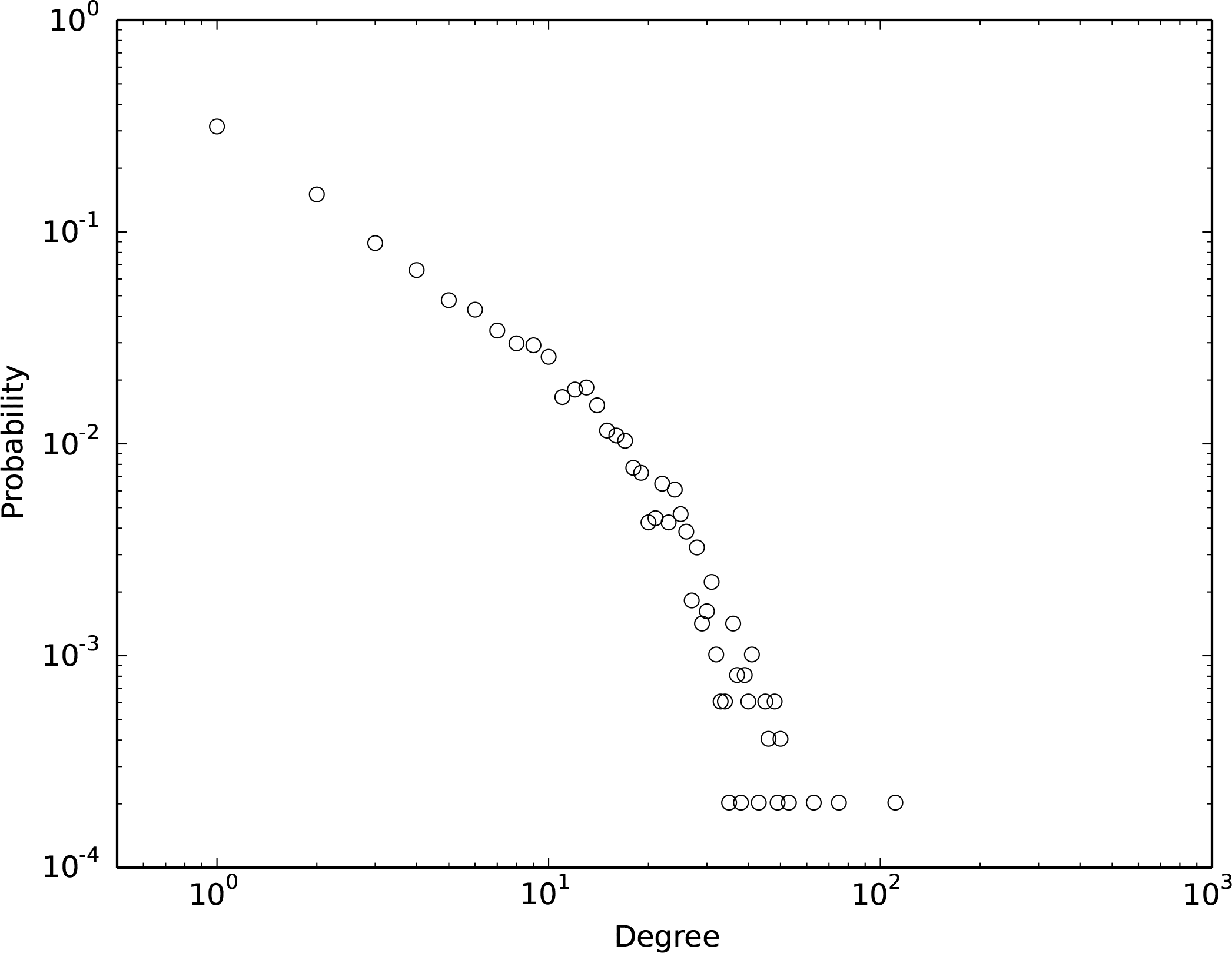
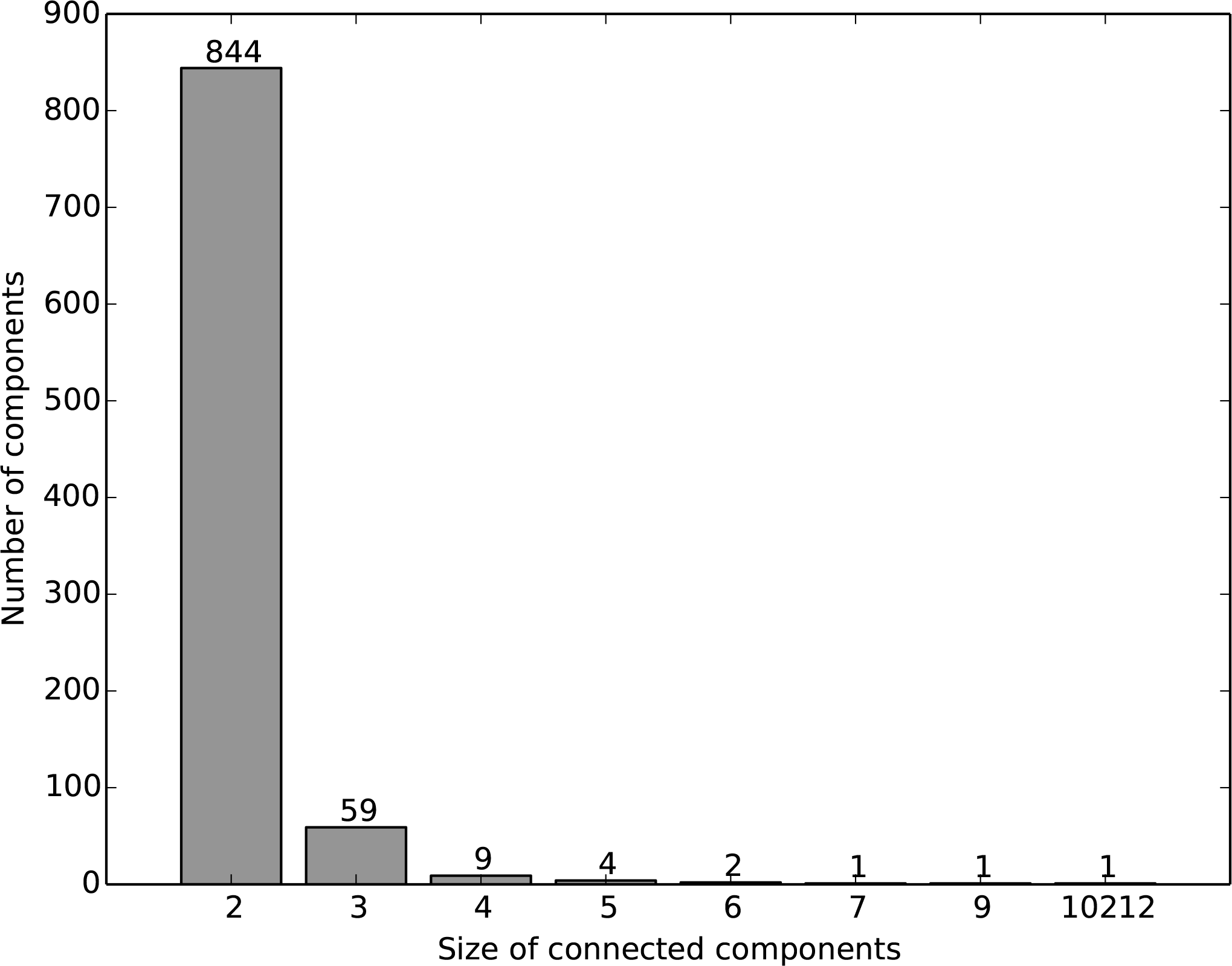
Degree distribution of a) diseases and b) genes in the bipartite disease-gene association network. The distributions are shown on a log-log scale.

**Fig 3.**
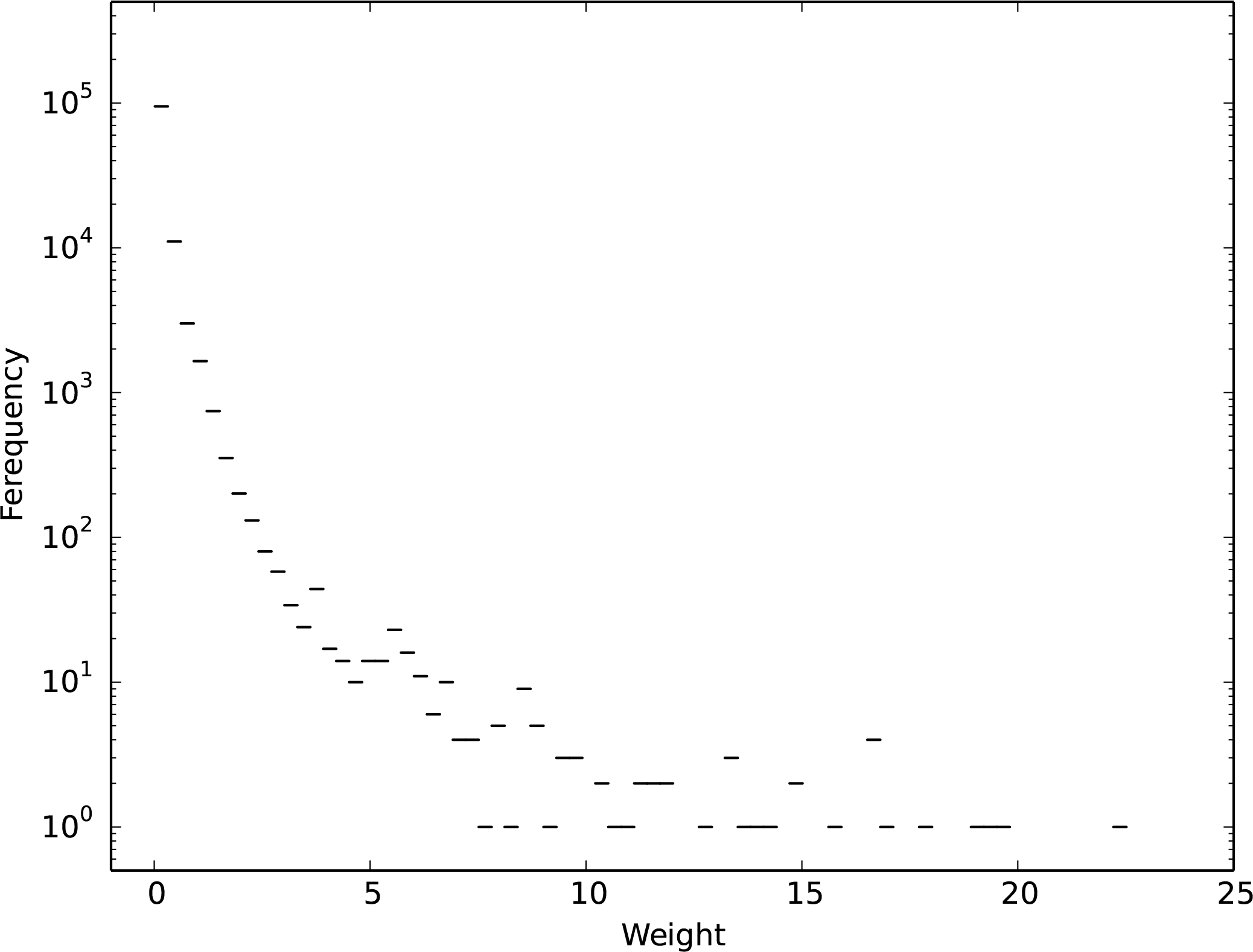
The size distribution of the connected components in the bipartite disease-gene network. The network has a single giant component with 10,212 vertices, and the majority of other connected components are of size two, i.e., consisting of only one disease and one gene.

#### Weighted Human Disease Network (WHDN)

We construct the weighted human disease network (WHDN) using the giant connected component of the bipartite disease-gene network. We use *D* and *G* to denote sets of 5,278 diseases and 4,934 genes respectively in the giant connected component. In the WHDN, an edge links two diseases *i* and *j* if they have at least one association gene in common, and the weight of the edge, *w*_*ij*_, is computed based on the number of shared association genes, as well as the strengths of those associations.

Such a weight definition is inspired by Newman’s study on scientific collaboration networks [14], where vertices are scientists and two scientists are connected by an unweighted edge if they have coauthored one or more scientific papers together. To define the strength of the tie between two connected scientists, two factors are considered. First, two scientists whose names appear on a paper together with many other coauthors know one another less well on average than two who are the sole authors of a paper. Thus, the collaborative ties are weighted inversely according to the number of coauthors of a paper. Second, authors who have written many papers together will know one another better on average than those who have written few papers together. Thus, all coauthored papers are added up to account for the tie strength of two scientists.

Here, similarly, first we consider that the correlation of two diseases through a gene is stronger when they are the sole associated diseases with this gene than when there are many other diseases associated with the same gene. Second, the correlation of two diseases is considered stronger when they share more genes through stronger associations than less genes or weaker associations. Thus, we extend Newman’s method to weighted graph and define the weight of edge W_ij_ between two diseases *i* and *j* as

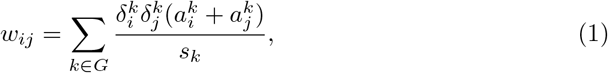

where 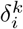 is one if disease *i* and gene *k* have a DGA, and zero otherwise. 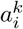 is the score of their DGA assessed by DisGeNET as discussed in the previous section, and *s*_*k*_ is the strength of gene *k* as a vertex in the bipartite disease-gene network, defined as the sum of the scores of the DGAs between gene *k* and its directly linked diseases,

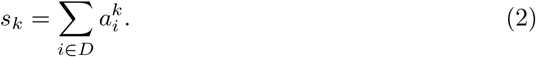

**Fig 4.**
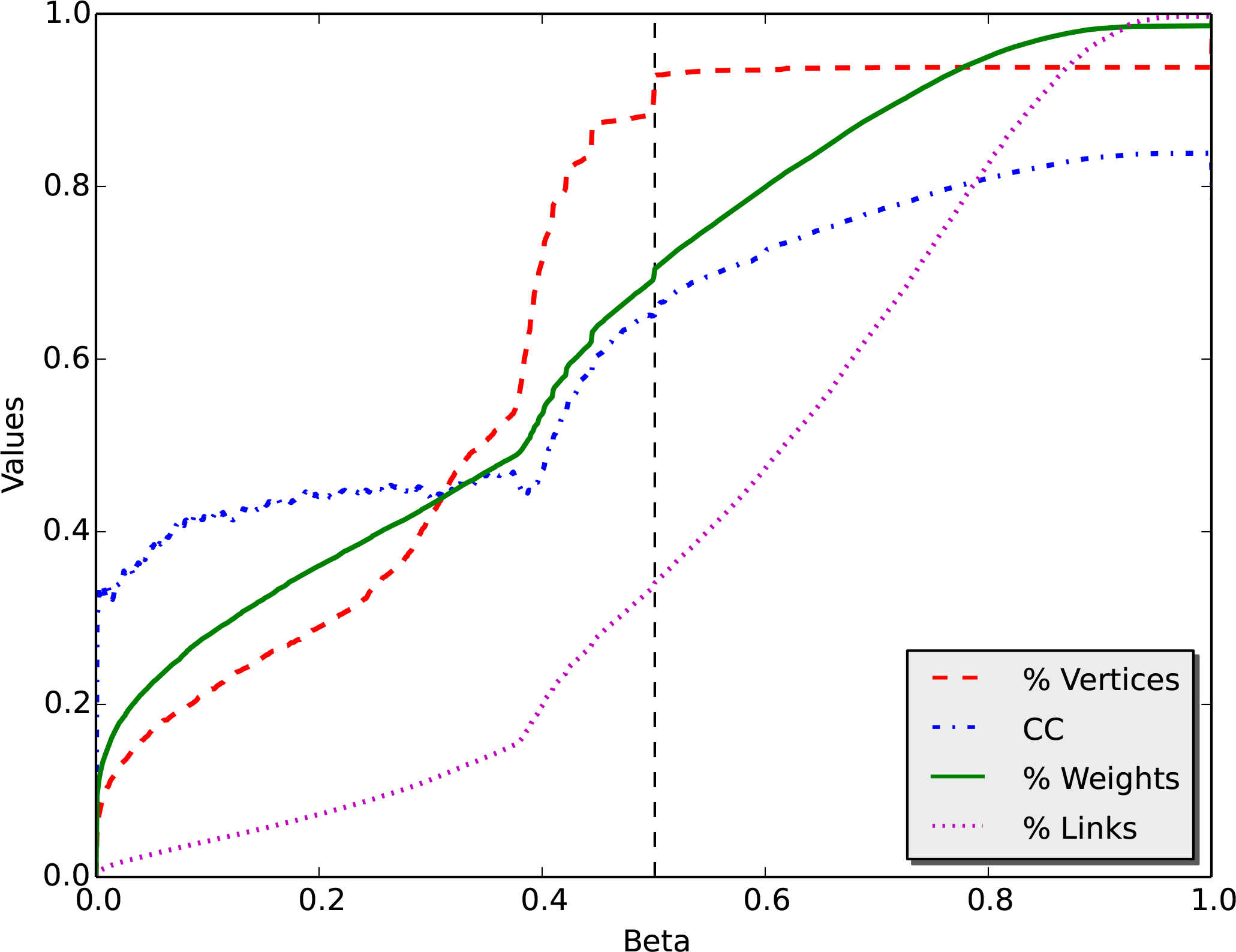
Distribution of edge weights in the WHDN. The weight of an edge quantifies the shared genetic background of two connected diseases. There are 112,324 edges in the graph with weights ranging from 0.0152 to 22.4506.

Such a weight definition indicates that the correlation strength of two diseases is weighted inversely according to the strengths of the genes they share, and is proportional to the total number of genes they share and the strengths of their DGAs.

For example, in Figure 1, the weight between diseases *contact dermatitis* (CD) and *white sponge nevus 1* (WSN1) is calculated as follows,

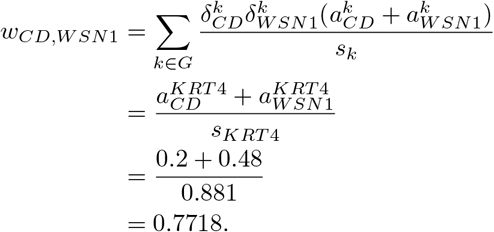

Note that the weight of two diseases can be greater than one when they share multiple genes. For example the weight between diseases WSN1 and *hereditary mucosal Leukokeratosis* (HML) is calculated as follows,

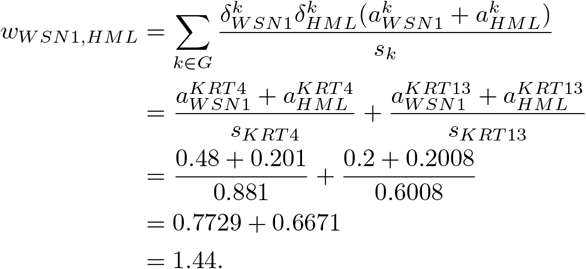

Since the WHDN is constructed using vertices from the giant component of the bipartite disease-gene association network, it only has a single connected component with all 5,278 vertices in the disease set *D*. Two vertices have an edge connecting them if the represented two diseases have at least one shared gene, and the edge weight is assessed as described above. The WHDN has 11,2324 edges and an average vertex degree of 42.56. That is, a disease correlates with on average 42.56 other diseases with varying strengths. Figure 4 depicts the distribution of all the edge weights in the WHDN. As we can see that a large number of edge weights are of small values and may not be particularly interesting for the subsequent analysis. Those weak edges not only add computational overhead to the network analysis, but also render the network difficult to interpret. Therefore, next we perform an edge reduction and only extract the most meaningful structure of the network.

#### The Multi-Scale Backbone of WHDN

The most straightforward strategy for network reduction is to use a global weight threshold and remove all links that have weights lower than the threshold. However, such a global thresholding strategy is somewhat arbitrary and may overlook the network information present below the cutoff scale. Here, to preserve the multi-scale backbone of the weighted human disease network (WHDN) while removing less relevant and meaningful edges we use a multi-scale filtering method proposed by Serrano *et al.* [43]. Such a multi-scale backbone exaction algorithm has been used to reduce the network size while preserving the meaningful structure of biological networks in multiple studies [32, 44, 45, 46].

First, the weight of edge linking vertex *i* with its neighbor *j* can be normalized as

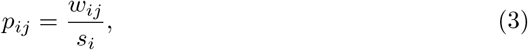

where *s*_*i*_ is the vertex strength, i.e., the sum of weights incident to vertex *i*, defined as

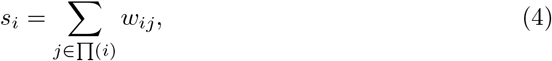

where Π(*i*) is the set of vertex *i*’s neighbors. Therefore, there are two different normalized values for a link *e*_*ij*_ using the strengths of its two end vertices *s*_*i*_ and *s*_*j*_ as the denominator.

Second, a null model is used to assess the expectation if the weights of links connecting to a particular vertex were distributed randomly. That is, the normalized weight *p_ij_* that corresponds to the link connecting to a certain vertex of degree *k* is produced by a random assignment from an uniform distribution. Thus the probability density function for the variable taking a particular value *x* is

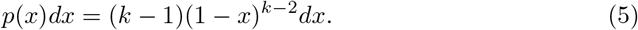

Then, to identify whether the probability, *β_ij_*, of link weight *p*_*ij*_ is compatible with the null model with a threshold *β* is given as

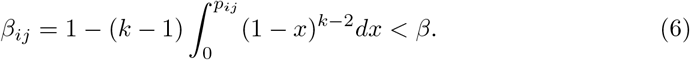

All links with computed *β*_*ij*_ lower than a given threshold *β* are preserved in the network. Note that each edge has two different values *β*_*ij*_ and *β*_*ji*_. For solving this problem, OR and AND rules can be used. Under the first rule, if either *β*_*ij*_ and *j* is lower than *β*, the link will be preserved. In the second case, an edge is preserved if both *β*_*ij*_ and *β_ji_* are lower than *β*. Darabos *et al.* [44] empirically found that the AND rule preserve the network features better than using the OR rule in the context of human phenotype networks. In this article, the AND rule is adopted to reduce the size of the network by removing the links which are less relevant.

To find the best cutoff for *β*, we calculate clustering coefficient, percentage of remaining vertices and links, and total weight of the networks after applying a *β* cutoff while *β* changes from 0 to 1. Figure 5 shows the results of network metrics as a function of *β* cutoffs. We choose a *β* cutoff when the clustering coefficient and the remaining vertices and weights are maximally preserved while as many links are removed as possible. Accordingly, the cutoff *β* = 0.501 can be determined, shown as the vertical dashed line in the figure.

After the backbone extraction, the WHDN has 4,898 vertices and 38,275 edges. Those vertices are no longer connected in a single component. Figure 6 shows the size distribution of its connected components. There is a giant component with 4,810 vertices and its degree distribution is shown in Figure 7. Again the degree distribution is heavy tailed and resembles a power-law relationship. The vertex *epilepsy* has the highest degree of 576. This giant component will be the focus for our next step analysis, e., measuring vertex importance in order to find the most central diseases in terms of correlating with other diseases.

**Fig 5.**
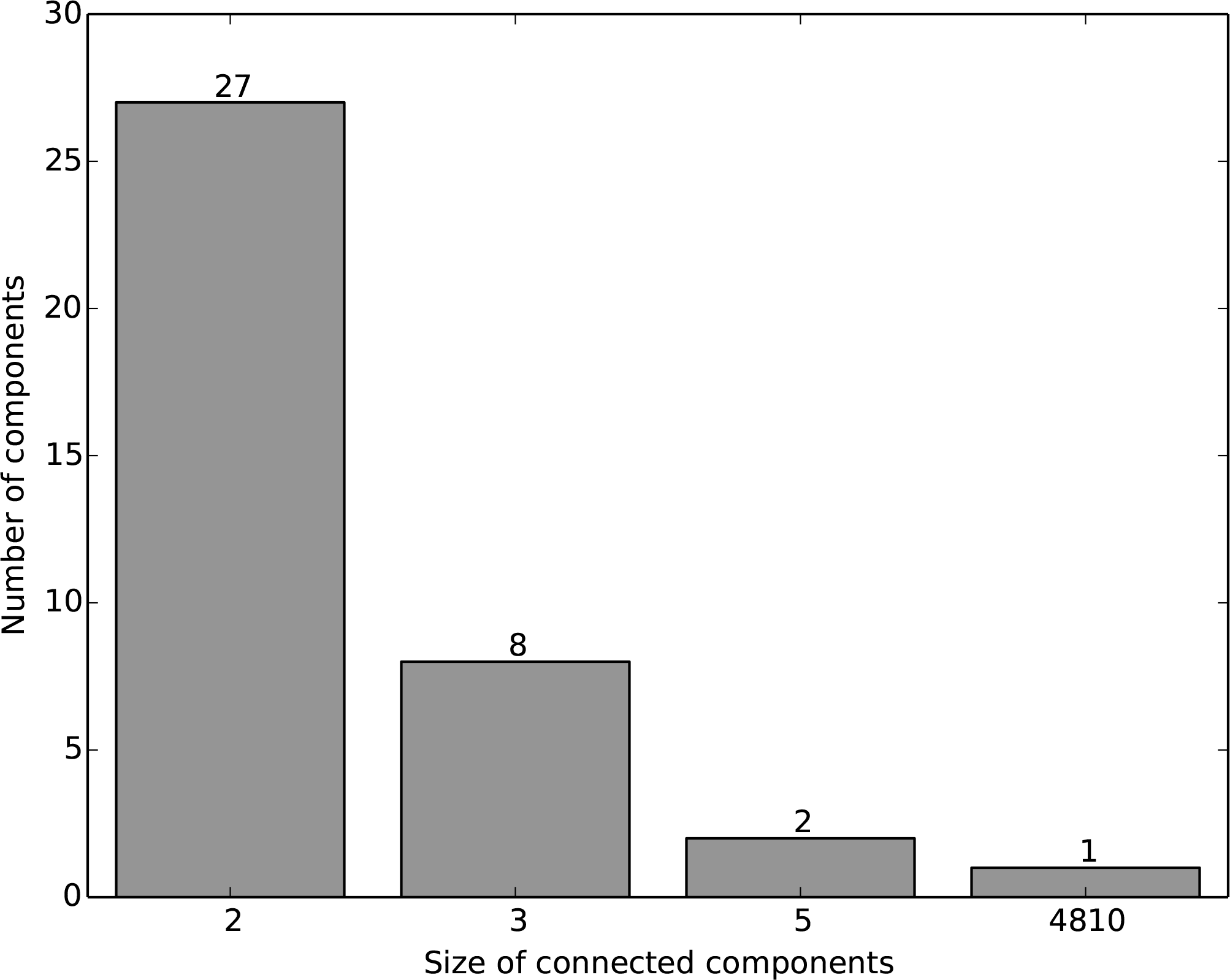
Choosing the *β* value. CC represents clustering coefficient, %Vertices is the percentage of remaining vertices, %Weights is the percentage of weights left after removing links, and %Links is the percentage of remaining links.

**Fig 6.**
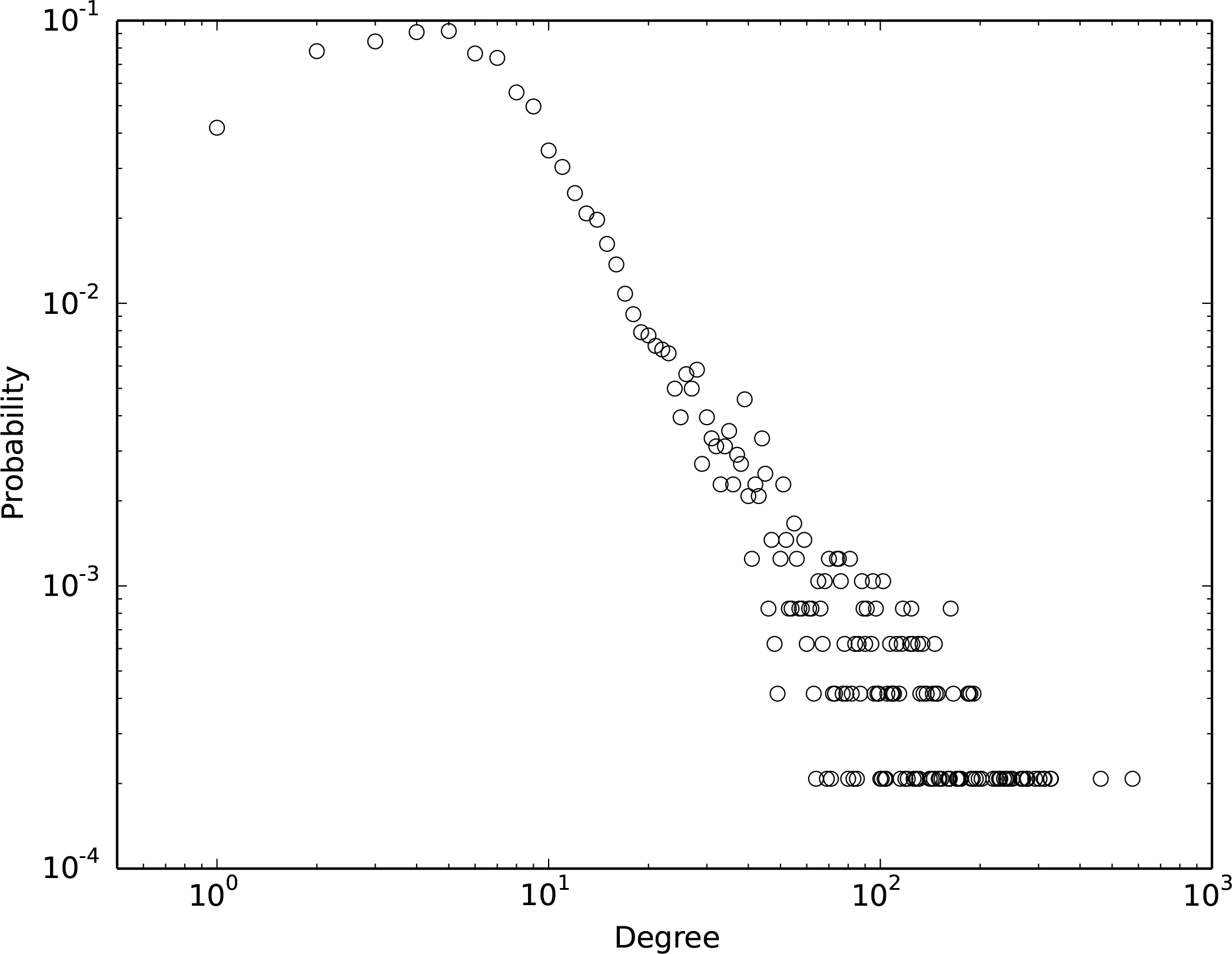
The size distribution of connected components in the extracted backbone of the WHDN. The network has a single giant component with 4,810 vertices.

**Fig 7.**
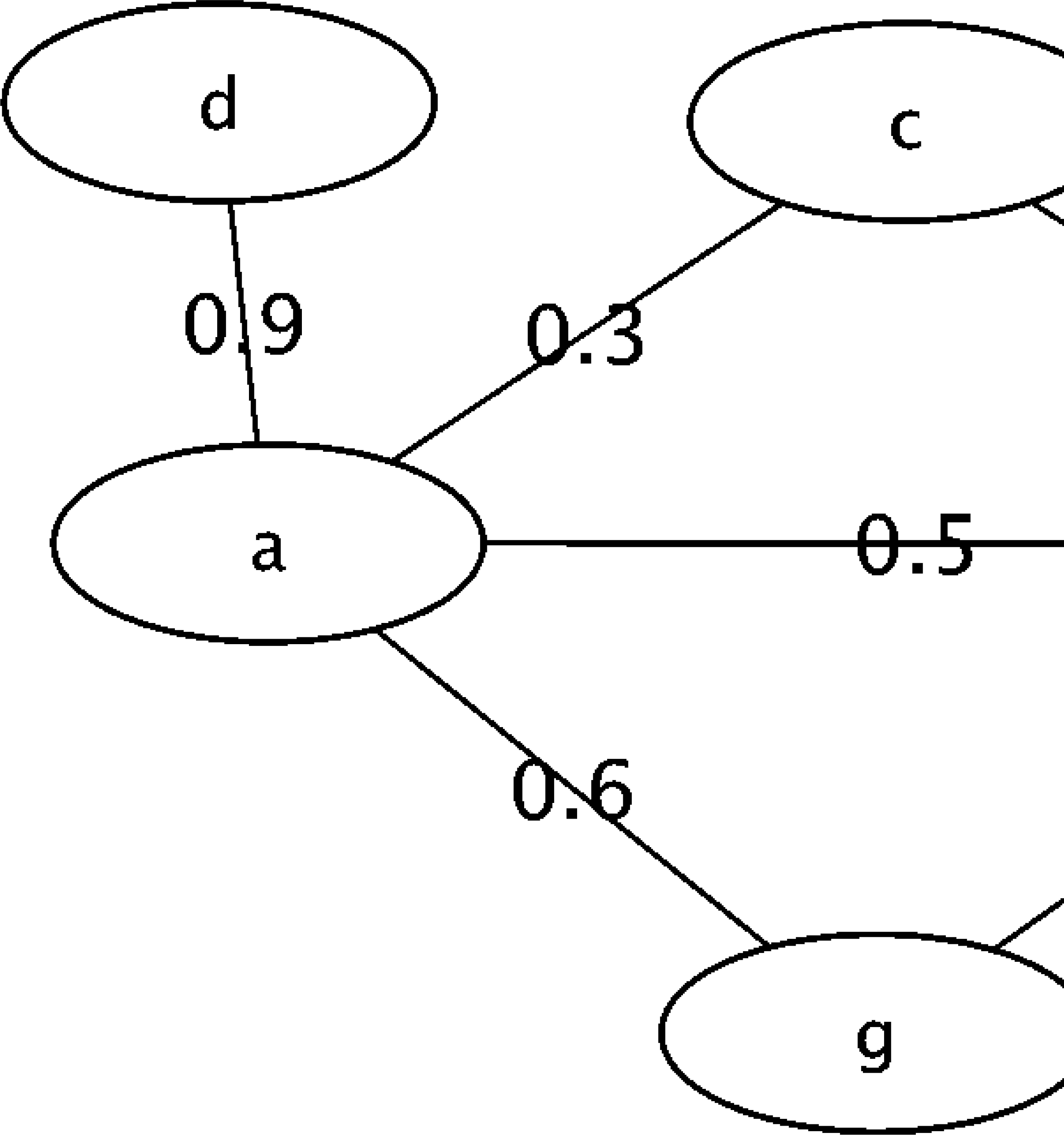
Degree distribution of vertices in the giant component of the extracted backbone of the WHDN. The distribution is shown on a log-log scale.

**Fig 8.**
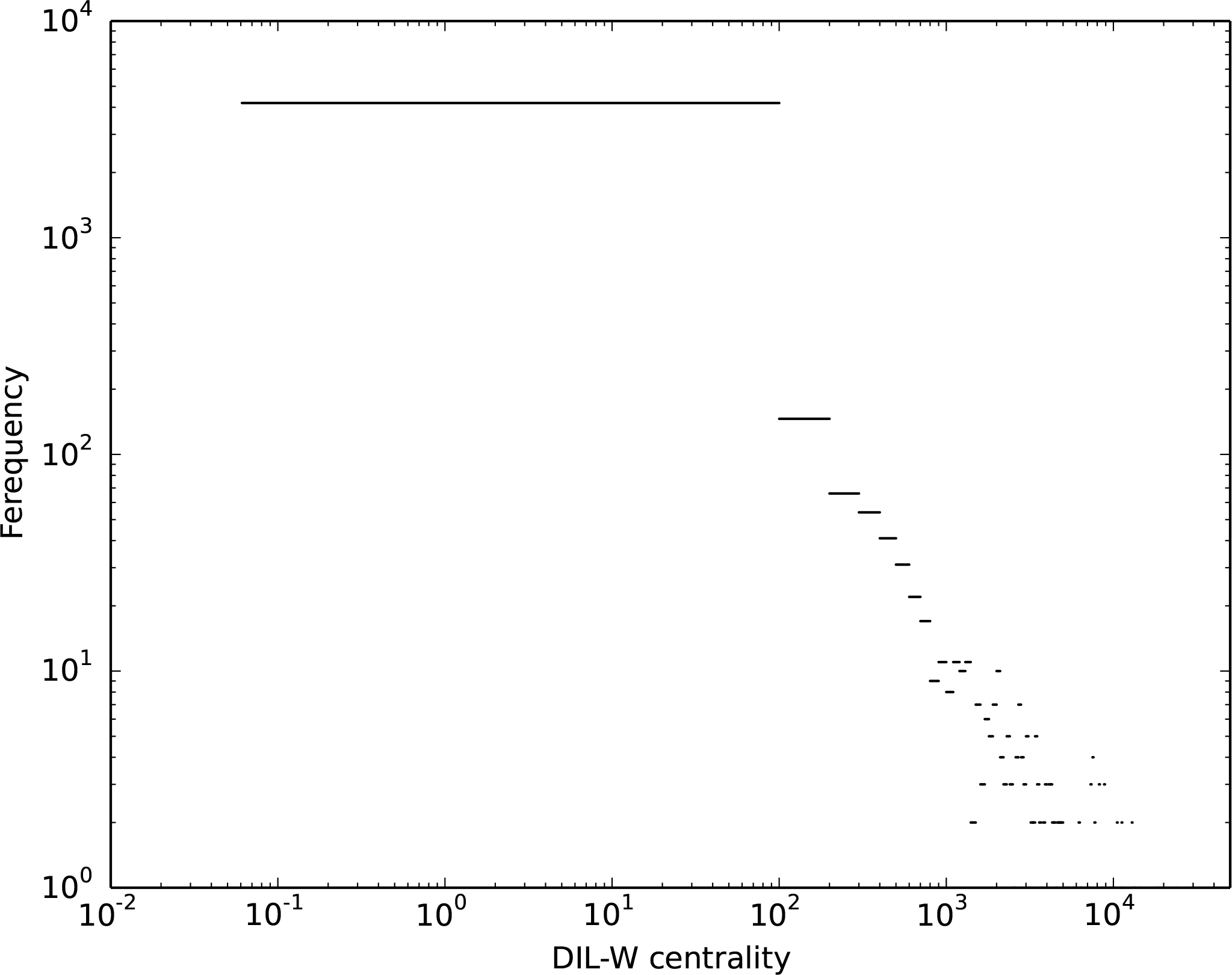
An example weighted graph.

### Measuring Vertex Importance in WHDN

Although various vertex centrality measures have been proposed in the literature, the quantification of the importance of a vertex in a network is often context-specific. For some networks, measuring degree may suffice since a vertex can be considered important when its number of neighbors is the sole criterion. For some networks, e.g., information communication networks, a vertex may be considered more important if its distances to all other vertices are short, then closeness centrality serves this purpose well. For our WHDN, a disease is considered important if it correlates with many other diseases (degree) as well as if the correlations are themselves very important (edge importance).

We propose a vertex importance measure for the weighted human disease network (WHDN) by extending a centrality measure for unweighted networks proposed by Liu *et al.* [47]. This measure assesses the centrality of a vertex based on both its degree and the importance of its incident links (DIL centrality). For its extension on weighted graphs, we name it the DIL-W centrality.

First, in the context of unweighted graph, the importance of a link *e_ij_* that connects vertex *v*_*i*_ and *v*_*j*_ can be calculated as follows:

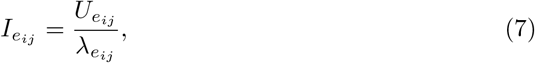

where 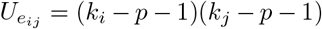 and 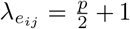. Following the convention, *k*_*i*_ and *k*_*j*_ are the degrees of vertex *v*_*i*_ and *v*_*j*_, respectively, and *p* is the number of triangles with one edge being *e*_*ij*_.

Subsequently, the contribution that vertex *v*_*i*_ makes to the importance of *e*_*ij*_ is computed as

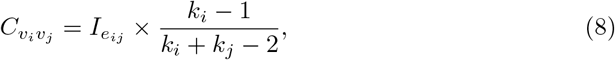

where *j* ∈ Γ_*i*_, and Γ_*i*_ is the neighborhood of vertex *i*.

Then, the DIL centrality of vertex *v*_*i*_ is calculated by combining both its degree and the importance of its incident links,

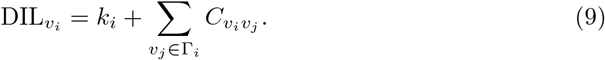

For weighted networks, we modify the computation of *U* in Equation (7) as

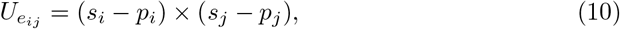

where *s*_*i*_ is the strength of vertex *v*_*i*_, calculated by Formula (4), and *p*_*i*_ is the weight sum of links incident to vertex *v*_*i*_ that form triangles with *e*_*ij*_. This follows the intuition that first an edge is considered more important when its two end vertices have higher strengths. Second, the importance of an edge is reduced when it has alternative two-hop paths connecting the same set of end vertices. Therefore, we subtract *p*_*i*_ from *s*_*i*_ in Equation (10).

We define λ for weighted graphs as

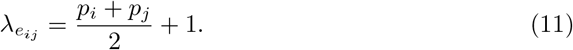

Finally, the importance of a vertex can be measured by

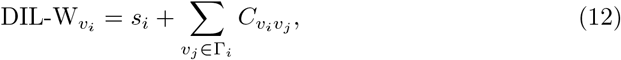

where 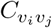 is defined as

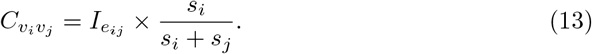

In the weighted graph given in Figure 8, vertex *a* has a higher strength but a lower degree than vertex *b*. We compute their DIL-W centralities and investigate which one is more central when both factors are considered.

First we have their strength values *s_a_* = 0.9 + 0.3 + 0.5 + 0.6 = 2.3, and *s*_*b*_ = 0.2 + 0.11 + 0.2 + 0.7 + 0.5 = 1.71. Their neighborhoods are Γ_*a*_ = {*b, c, d, g*} and Γ_*b*_ = {*a, c, e, f, g*}. For vertex *a*,

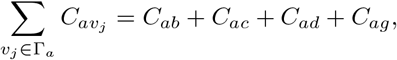

where

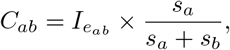

and

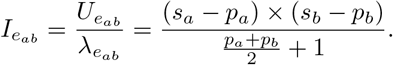

We have

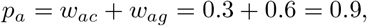

and

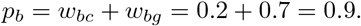

So

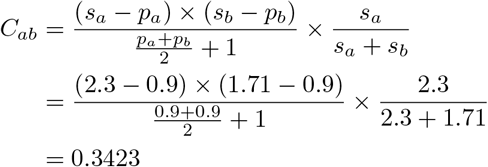

We can also have

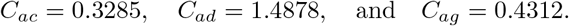

Then

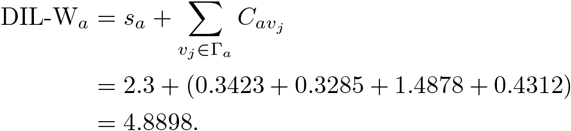

**Fig 9.**
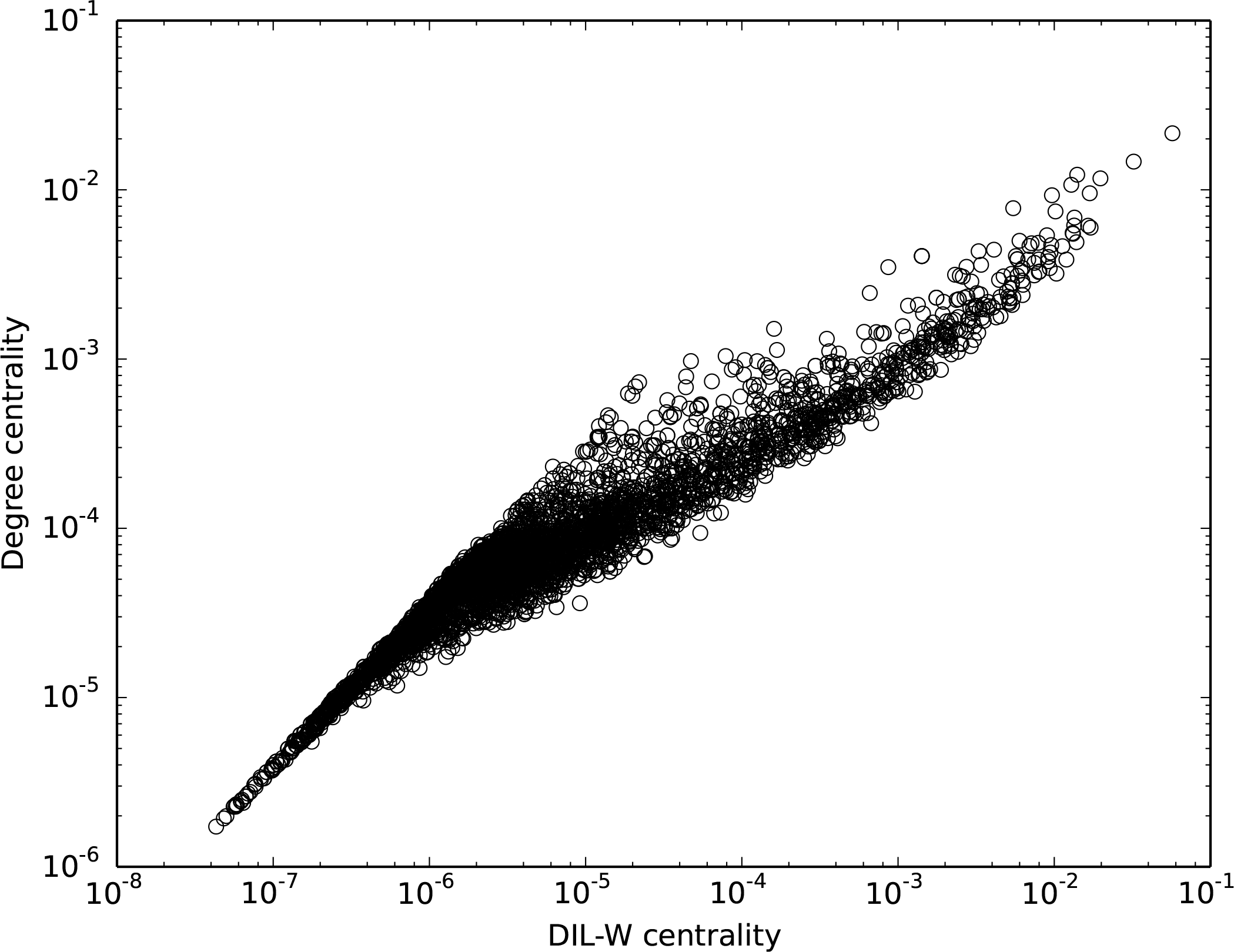
Distribution of DIL-W centrality in the giant component of the WHDN on a log-log scale.

Similarly, we can compute the DIL-W centrality of vertex *b* DIL-W_*b*_ = 2.8916. Therefore, based on both the degree and importance of incident edges, vertex *a* is considered more important than vertex *b*.

We apply the DIL-W centrality measurement to the giant component of the backbone of WHDN, the distribution is shown in Figure 9. The DIL-W scores have a high dynamic range, from 0.0610 to 80688.1129. The majority of the vertices have low scores and a few number of vertices can have scores that are greater by orders of magnitude.

**Fig 10.**
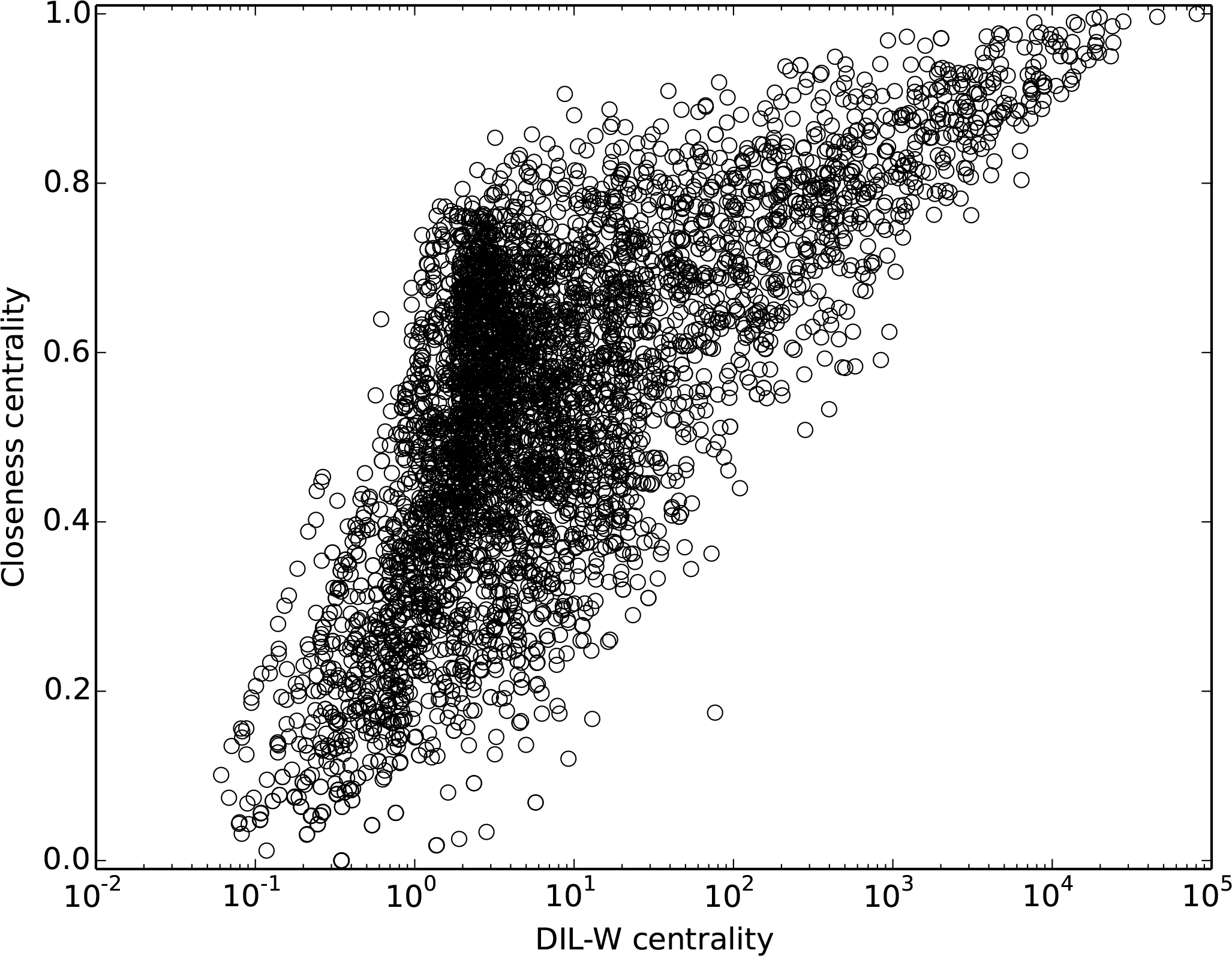
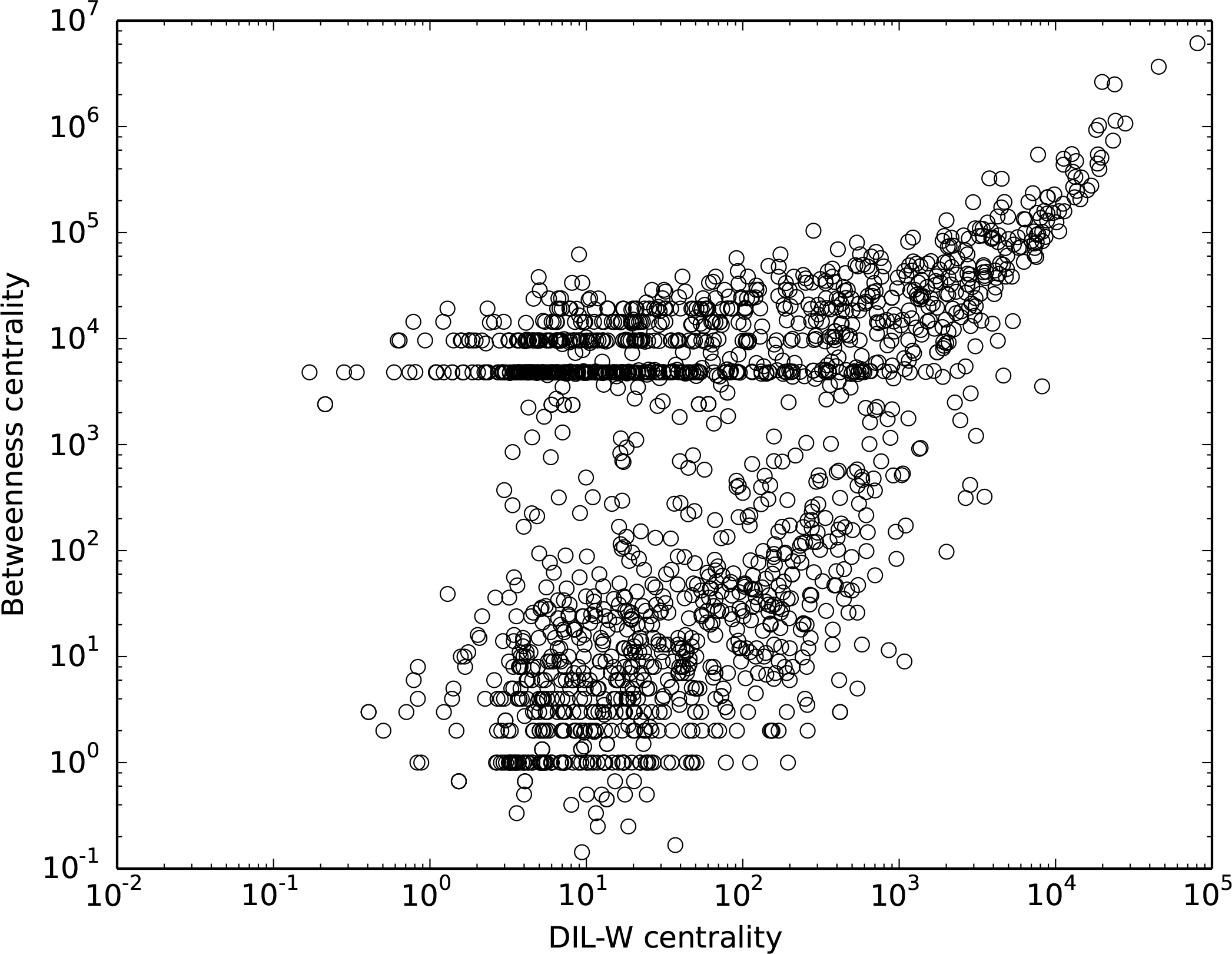
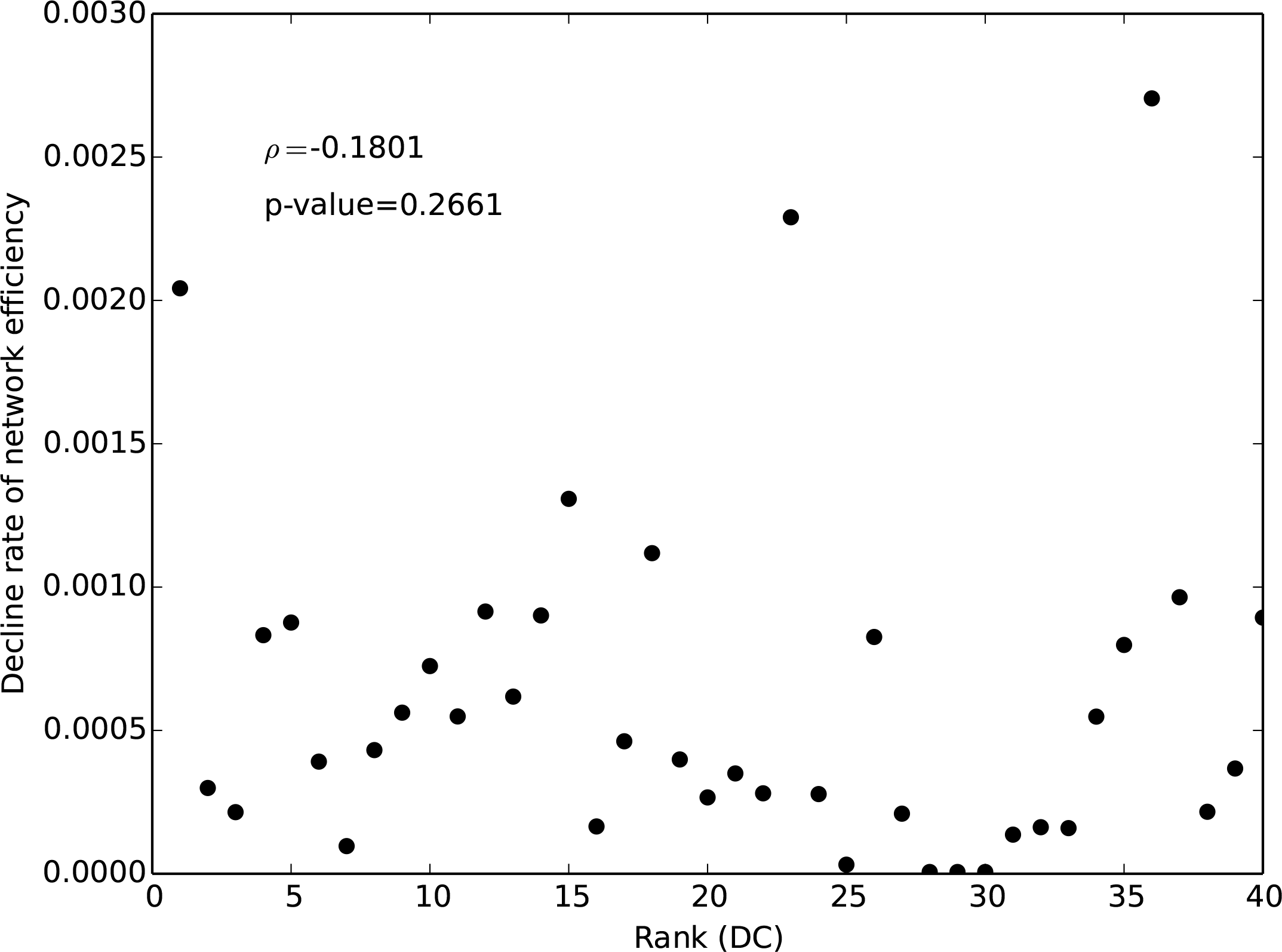
Correlation of DIL-W scores with a) degree centrality, b) closeness centrality, and c) betweenness centrality in the WHDN.

### Comparison and Evaluation

We compare our DIL-W measurement with three most commonly used centralities, i.e., degree, closeness, and betweenness, when applied to the giant component of the backbone of WHDN. For weighted graphs, degree centrality is calculated as vertex strength given by Equation (4). Closeness and betweenness are shortest-path-based centralities. Shortest path computation can be extended for weighted graph as follows,

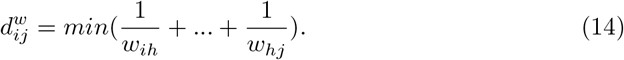

Here 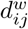 denotes the weighted distance between vertex *i* and *j*, and *w*_*ih*_ is the weight of the edge linking vertex *i* and *h*. Since in our WHDN edge weight suggests strength, the distance between two vertices is the minimum sum of the inverse of edge weight along the path connecting them. Once the weighted distance is defined, closeness and betweenness can be calculated by their original definitions.

Figure 10 shows the correlation of DIL-W scores with a) degree, b) closeness, and c) betweenness centralities. As we can see, there is a positive correlation between DIL-W measure and all other three vertex centrality measures. The Spearman’s rank correlation coefficient is 0.672 comparing DIL-W with closeness, is 0.71 comparing DIL-W with betweenness, and is 0.947 comparing DIL-W with degree.

To evaluate our new vertex importance quantification method, DIL-W, we measure the network efficiency before and after we remove the most important vertices in the WHDN. In the context of the WHDN, the network efficiency indicates the extend to which the original connectivity of the network is maintained. We calculate the decline rate of network efficiency after removing *m* top-rank vertices. The network efficiency [48] is computed based on the connectivity of a network. A higher connectivity suggests a higher network efficiency. The network efficiency is defined by

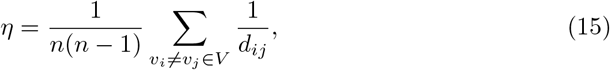

where *n* is the total number of vertices in the network, *V* is the vertex set, and *d*_*ij*_ is the weighted distance between vertex *v*_*i*_ and *v*_*j*_. Thus, the decline rate of the network efficiency is calculated as

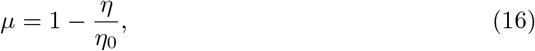

where *η*_0_ is the efficiency of the original network, and *η* is the network efficiency after some vertices are removed.

**Fig 11.**
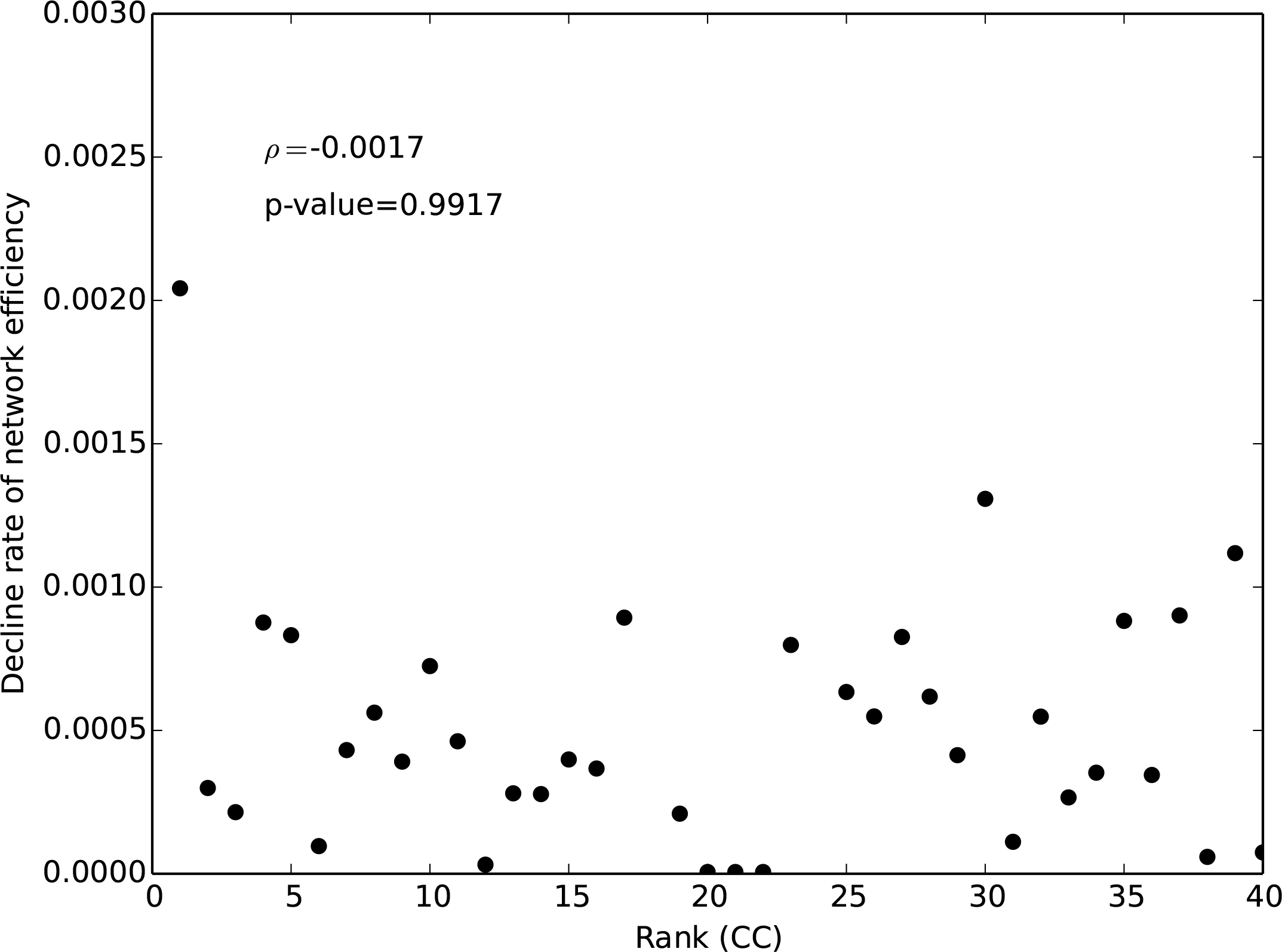
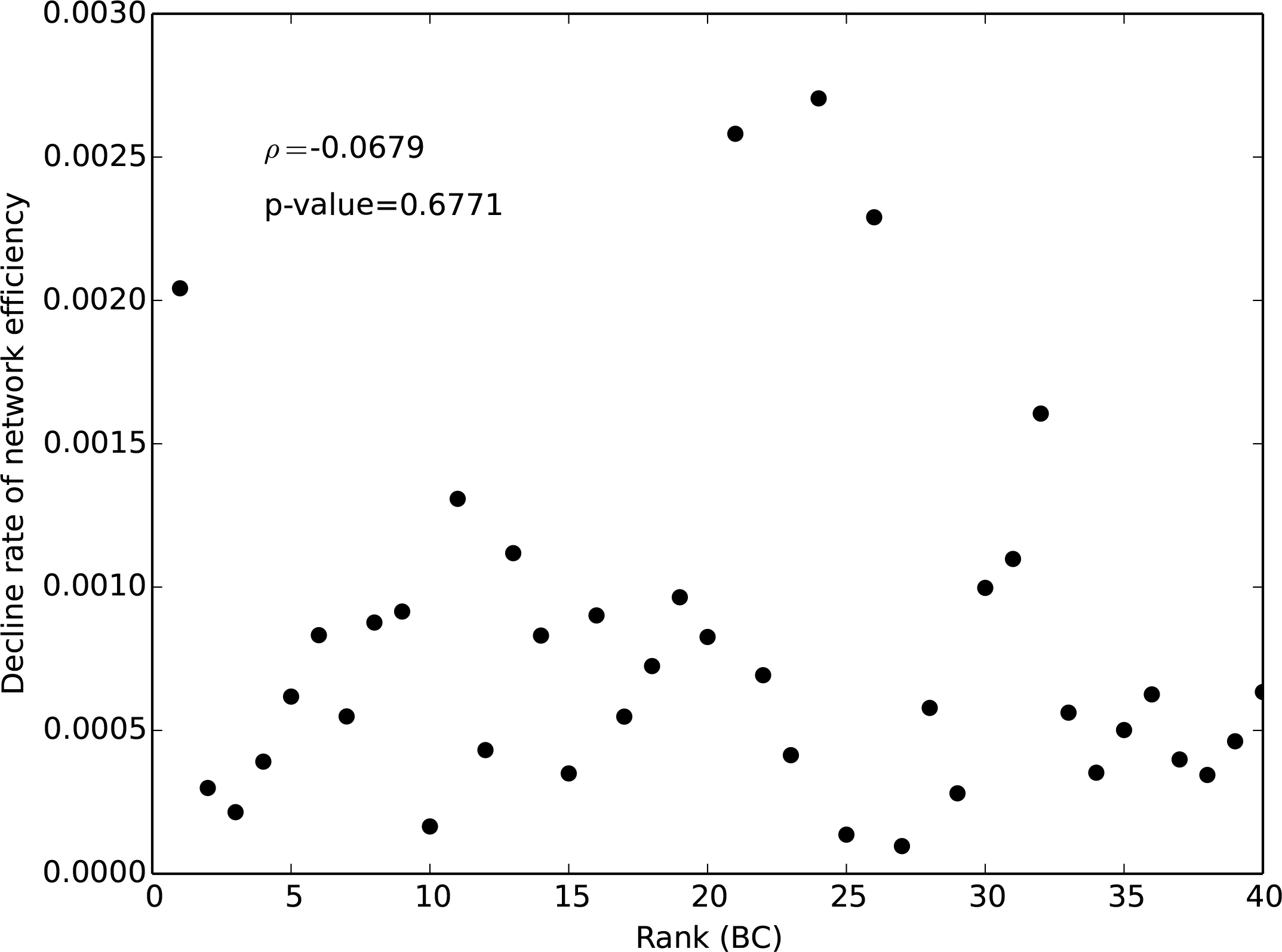
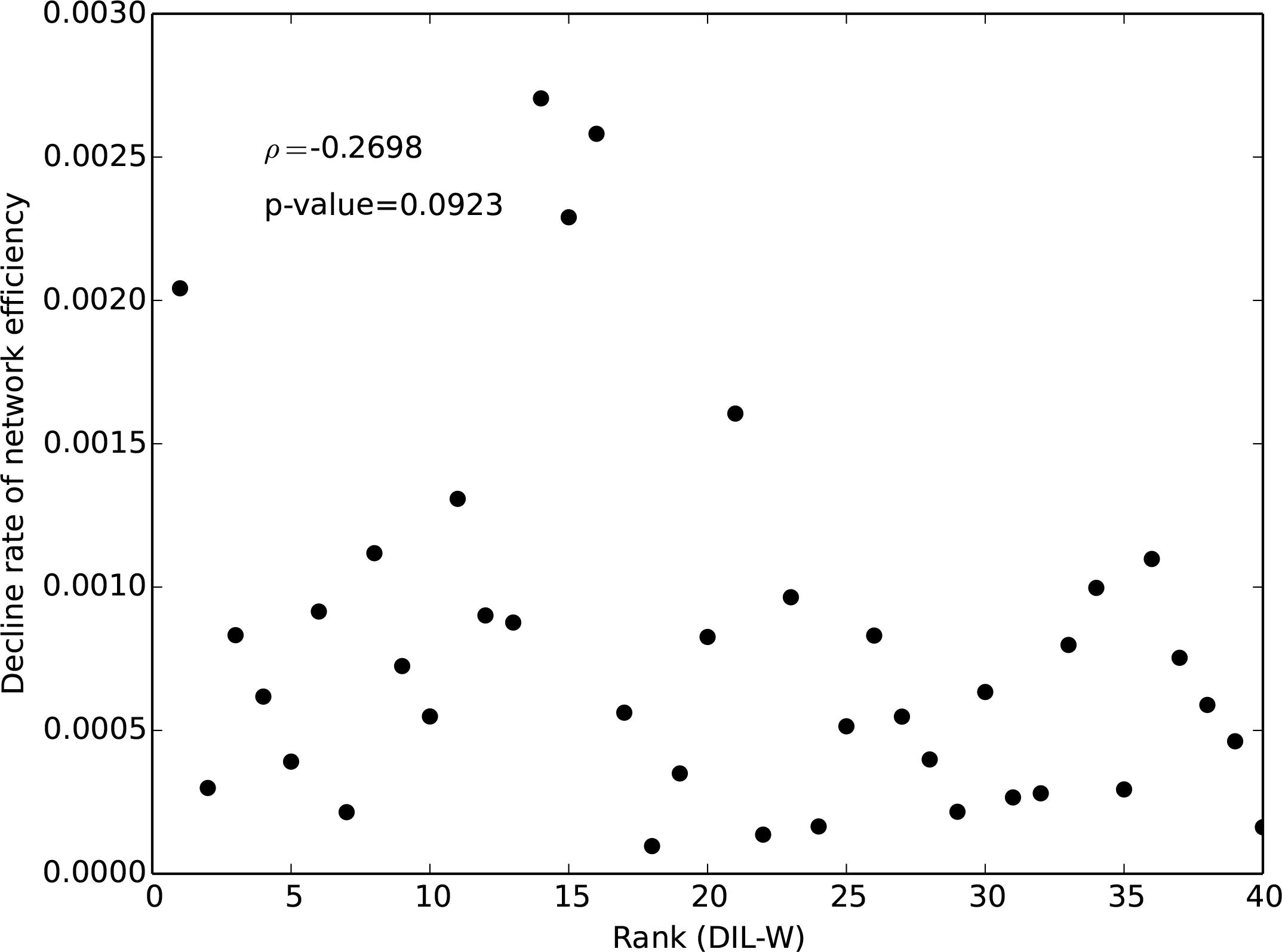
Decline rate of network efficiency after removing a single vertex ranked by a) degree centrality (DC), b) closeness centrality (CC), c) betweenness centrality (BC), and d) DIL-W.

**Fig 12.**
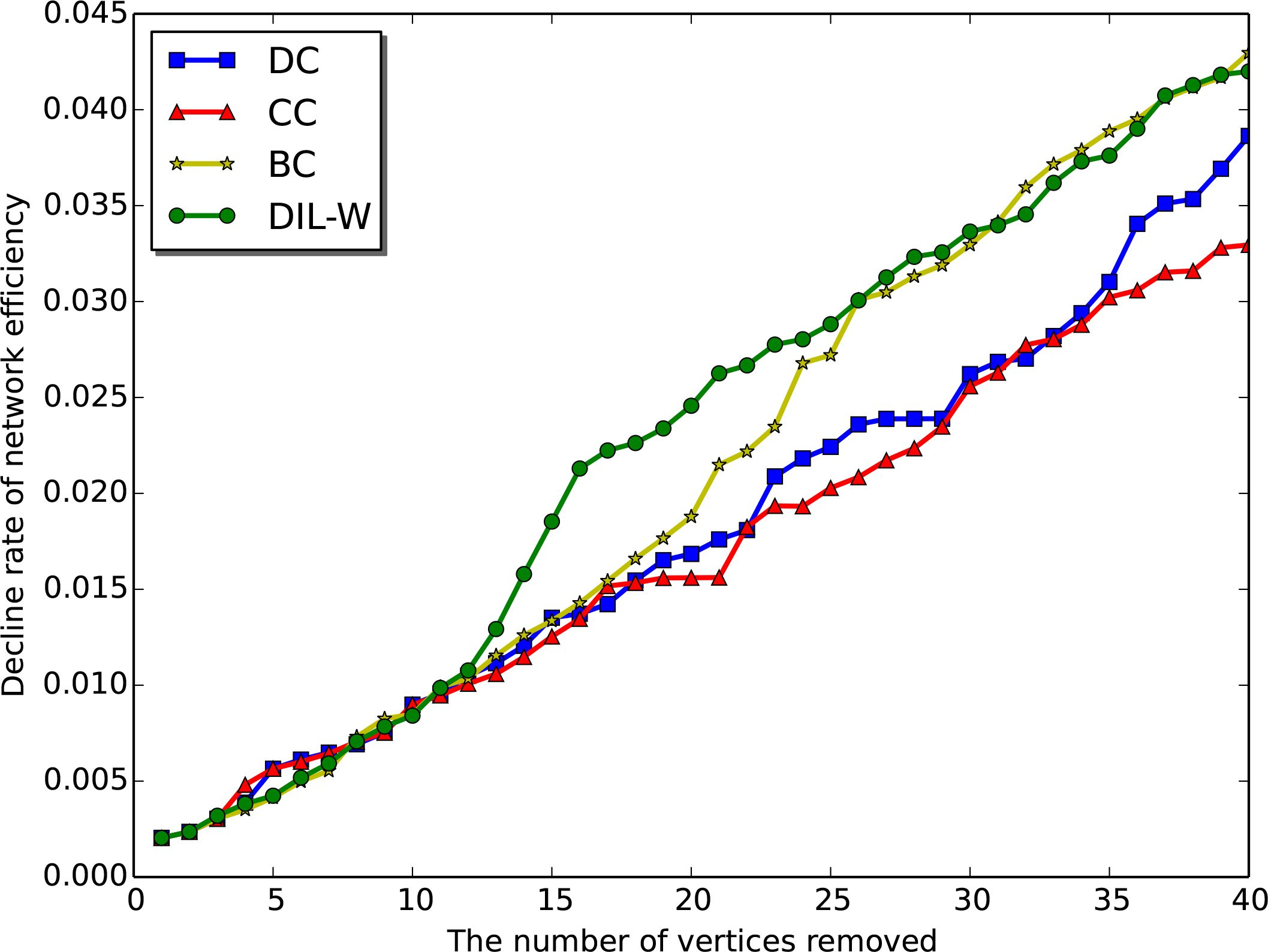
The decline rate of the network efficiency as a function of removing the top m vertices ranked by degree centrality (DC), closeness centrality (CC), betweenness centrality (BC), and DIL-W.

When a more importance vertex is removed, we expect to see a greater decline rate of the network efficiency. Thus we can use *μ* as a indicator for the actual impact of removing a vertex in the network. Figure 11 shows the decline rate of the network efficiency when we remove each of the top 40 vertices ranked by a) degree (DC), b) closeness (CC), *c*) betweenness (BC), and d) DIL-W. Further removal of top ranked vertices could be investigated but was not included in the current study given the high computational demand. As shown in the figure, we do not observe a monotonic relationship across all four centrality methods. However, the correlation analysis shows that our method, DIL-W, has a slighter stronger negative correlation between the decline rate and the rank of the removed vertex than the other three. The Spearman’s rank correlation coefficient, *ρ*, for degree, closeness, and betweenness is −0.1801, −0.0017, and −0.0679, respectively. In comparison, DIL-W has a negative correlation coefficient −0.2698.

We also consider removing all *m* top-rank vertices at once and see how this accumulative removal affects the efficiency of the network. Figure 12 shows the decline rate of the network efficiency after removing all top *m* vertices ranked by different centrality measures. The graph shows that the proposed method, DIL-W, has the highest decline rate of network efficiency for 57.5% of the data points, while betweenness, closeness, and degree have 27.5%, 10%, and 5%, respectively. This suggests that DIL-W is able to select a set of more important vertices comparing with the other three centrality measures. As seen in Figure 12, the four methods are very comparable until the top 11 diseases are removed from the network. Then DIL-W has a significant higher network efficiency decline rate than the rest. Betweenness centrality catches up around point 30 and becomes very comparable afterwards.

Table 1 shows the top 30 diseases ranked by our DIL-W method, their degrees, and their neighbors that have the strongest correlations (i.e., edge weights). References that support the known comorbidity of the disease pairs are also given.

**Table 1.** The 30 top-ranked diseases by DIL-W and their most correlated dieases

| Rank | Disease | Degree | The most correlated disease | Ref. |
| --- | --- | --- | --- | --- |
| 1 | Epilepsy | 576 | Pediatric failure to thrive | – |
| 2 | Pediatric failure to thrive | 462 | Epilepsy | – |
| 3 | Sensorineural hearing loss (disorder) | 313 | Retinitis pigmentosa | [54] |
| 4 | Anemia | 327 | Pediatric failure to thrive | [50] |
| 5 | Obesity | 268 | Retinitis Pigmentosa | [49] |
| 6 | Osteoporosis | 326 | Osteopenia | [55] |
| 7 | Nystagmus | 276 | Epilepsy | [56] |
| 8 | Liver cirrhosis | 278 | Chemical and drug induced liver injury | [51] |
| 9 | Low vision | 270 | Nystagmus | [57] |
| 10 | Heart failure | 311 | Obesity | [58] |
| 11 | Muscle degeneration | 277 | Amyotrophic lateral sclerosis | [59] |

**Table 1.** The 30 top-ranked diseases by DIL-W and their most correlated diseases
| Rank | Disease | Degree | The most correlated disease | Ref. |
| --- | --- | --- | --- | --- |
| 12 | Diabetes mellitus, non-insulin-dependent | 245 | Obesity | [60] |
| 13 | Strabismus | 293 | Epilepsy | [61] |
| 14 | Exophthalmos | 302 | Strabismus | [62] |
| 15 | Myopia | 266 | Sensorineural hearing loss (disorder) | [63] |
| 16 | Degenerative polyarthritis | 239 | Rheumatoid arthritis | [64] |
| 17 | Cerebral atrophy | 267 | Epilepsy | [65] |
| 18 | Optic atrophy | 236 | Nystagmus | – |
| 19 | Rheumatoid arthritis | 188 | Lupus erythematosus, systemic | [66] |
| 20 | Hydrocephalus | 250 | Epilepsy | [67] |
| 21 | Alopecia | 241 | Dystrophia unguium | – |
| 22 | Myocardial ischemia | 166 | Obesity | – |
| 23 | Myocardial infarction | 228 | Coronary artery disease | [68] |
| 24 | Chemical and drug induced liver injury | 174 | Cholestasis | [69] |
| 25 | Asthma | 198 | Dermatitis, atopic | [70] |
| 26 | Endometriosis | 135 | Obesity | [71] |
| 27 | Hypertrophic cardiomyopathy | 187 | Pediatric failure to thrive | [72] |
| 28 | Conductive hearing loss | 163 | Sensorineural hearing loss (disorder) | [73] |
| 29 | Brain ischemia | 191 | Diabetes mellitus, non-insulin-dependent | – |
| 30 | Gastroesophageal reflux disease | 190 | Epilepsy | [74] |

## Discussion

In this article, we use a network-based analysis to identify important human diseases that share genetic background with many other diseases through strong associations. We collect a large number of known disease-gene associations (DGAs) using DisGeNET in order to construct a bipartite disease-gene network. Subsequently, a weighted human disease network (WHDN) is built by connecting pairs of diseases that share associated genes and the edge weights reflect the number of genes they share as well as the strength of the DGAs. Then we propose a new vertex centrality measure DIL-W that considers both the degree of a vertex and the importance of its incident edges in weighted graphs. Upon application to the WHDN, DIL-W is shown to outperform three commonly used centrality measures, degree, closeness and betweenness, and has identified top diseases including *epilepsy, anemia*, and *obesity.*

Table 1 shows the degree in the WHDN and the most correlated disease of those 30 top-rank diseases. We are also able to find previous publications that verify almost all the correlations of those pairs of diseases, shown as references in the table. Besides some very well-known correlations such as *heart failure* - *obesity* and *diabetes* - *obesity*, the table also reports some less known but interesting correlations. For instance, Savin [49] showed that *atypical retinitis pigmentosa* is correlated with *obesity*. Moreover, the correlation between *anemia* and *pediatric failure to thrive* had not been reported in the literature until recently Dimmock *et al*. [50] suggested *anemia* as one of the novel causes of *failure to thrive* in children. Zimmerman [51] studied the cause of different types of *cirrhosis* resulting from different drug-induced injuries. This supports our finding on the correlation between *cirrhosis* and *chemical and drug induced liver injury*.

The disease-gene associations come from DisGeNet [42] only. While this is a valuable resource, it is merely one of the many databases that have disease gene information (including Jensen Lab’s DISEASES [52] and DiseaseConnect [53] databases), all of which have their own disease association scoring convention. The alternative databases will be explored in our future study.

## Conclusion

Apart from many existing related work, in this article, we construct a *weighted* human disease network (WHDN) and propose a new centrality measure DIL-W designed specifically for the WHDN. Our network-based analysis methods are shown to be able to identify more important diseases comparing to degree, closeness and betweenness centralities. The identified disease-disease correlations include previous knowledge supported by published literature as well as less known and novel correlations that can be valuable for future studies. Our understanding of human diseases is still largely unclear and the disease-gene associations are far from being complete. Future studies could explore the utilization of multiple types of data and more powerful computational tools to better cluster and categorize human diseases and to predict new genes and other factors that can explain diseases.

## Acknowledgments

This research was supported by the Discovery Grant from the Natural Sciences and Engineering Research Council of Canada (NSERC) RGPIN-2016-04699 to TH. The computation was feasible with the help from the IBM HPC cluster at the Center for Health Informatics & Analytics (CHIA), Faculty of Medicine, Memorial University.

